# Advanced Feedback Enhances Sensorimotor Adaptation

**DOI:** 10.1101/2022.09.14.508027

**Authors:** Tianhe Wang, Guy Avraham, Jonathan S. Tsay, Tanvi Thummala, Richard B. Ivry

**Author notes:** Corresponding authors: Tianhe Wang.

## Abstract

It is widely recognized that sensorimotor learning is enhanced when the feedback is provided throughout the movement compared to when it is provided at the end of the movement. However, the source of this advantage is unclear: Continuous feedback is more ecological, dynamic, and available earlier than endpoint feedback. Here we assess the relative merits of these factors using a method that allows us to manipulate feedback timing independent of actual hand position. By manipulating the onset time of ‘endpoint’ feedback, we found that adaptation was modulated in a non-monotonic manner, with the peak of the function occurring in advance of the hand reaching the target. Moreover, at this optimal time, learning was of similar magnitude as that observed with continuous feedback. By varying movement duration, we demonstrate that this optimal time occurs at a relatively fixed time after movement onset, an interval we hypothesize corresponds to when the comparison of the sensory prediction and feedback generates the strongest error signal.

## Introduction

Implicit adaptation ensures that the sensorimotor system remains exquisitely calibrated in the face of a variable environment and fluctuations in the internal state of the agent. This process occurs automatically in response to sensory prediction error, the mismatch between the expected sensory consequences of a movement and the actual feedback^1–3^. A common way to examine constraints on sensorimotor adaptation is to manipulate the visual error. For example, by occluding the arm and providing cursor feedback, a polar transformation (e.g., visuomotor rotation) can be used to introduce a discrepancy between the actual and perceived position of the hand. This discrepancy serves as an error signal that is used to recalibrate the sensorimotor system to minimize future errors when a similar action is produced.

In studies of visuomotor adaptation, two types of visual feedback are typically used: Continuous feedback where the cursor is visible throughout the movement and thus provides feedback of the movement trajectory, and endpoint feedback where the cursor is only presented when the hand reached its terminal position or at the radial distance of the target (Fig. 1c). It is well-established that adaptation in response to endpoint feedback is attenuated compared to adaptation in response to continuous feedback^4–6^. This effect is especially pronounced in measures of implicit adaptation, with little difference between online and endpoint feedback on measures reflective of strategic changes in performance^7^. Moreover, the efficacy of endpoint feedback is constrained by timing. Specifically, when the presentation of the feedback cursor is delayed relative to the hand movement, adaptation is markedly attenuated^8–12^. Indeed, delaying endpoint feedback by just 100ms can produce a dramatic reduction in the magnitude of implicit adaptation^13,14^.

**Figure 1.**
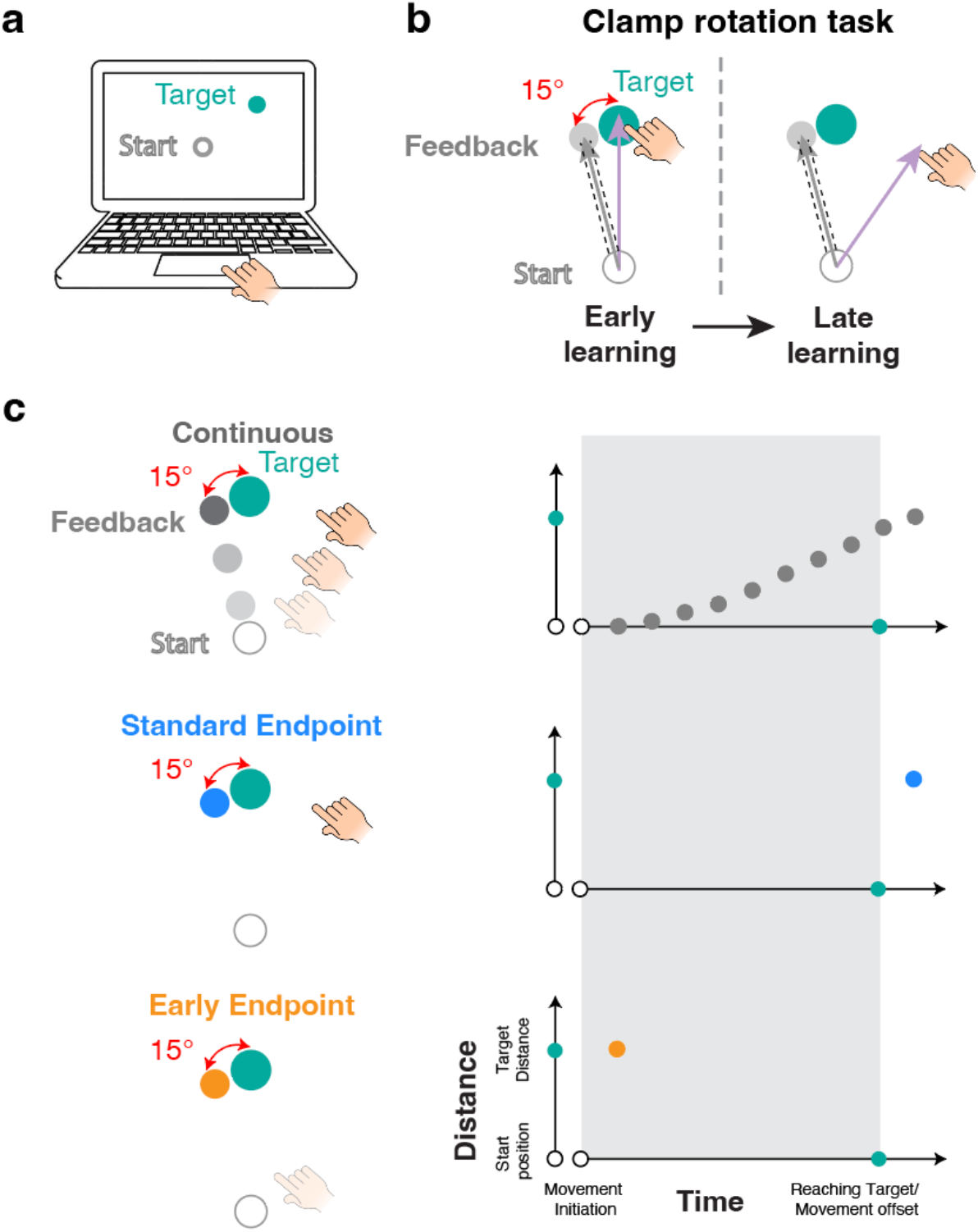
Experimental setup, task, and feedback conditions. **A**) Schematic of the web-based experimental setup, depicting the start location (white circle), the cursor (white dot), and a target (cyan dot). **B**) For task-irrelevant clamped feedback, the angular position of the feedback cursor is rotated by 15° with respect to the target, regardless of the heading direction of the hand. **C)** Three types of clamped feedback are illustrated in task space (left) and as a function showing the position and time of the feedback (right). Each dot represents one refresh cycle on the computer display monitor (16.7ms for most of the monitors). Continuous feedback is presented throughout the movement, following the radial distance of the hand from the start location to the target. Standard-endpoint feedback is presented for one refresh cycle after the hand reaches the target distance (corresponding to the last frame of the continuous feedback condition). Early-endpoint feedback is presented for one refresh cycle at the radial distance of the target when movement initiation is detected. The gray area indicates the interval between movement onset and when the radial distance of the movement reaches the target amplitude. Due to hardware limitations, there is a delay between detection of the hand movement and the onset of the feedback (relevant for continuous and early endpoint feedback), and a delay between when the hand reaches the target amplitude and presentation (relevant for standard endpoint feedback).

At present, it remains unclear why continuous feedback is advantageous relative to endpoint feedback. There are many notable differences between these two modes. First, by definition, continuous feedback is dynamic and endpoint feedback is static. Given the dynamic nature of most human sensorimotor skills, the nervous system may be more responsive to the ecological nature of continuous feedback. Second, continuous feedback provides a continuous stream of spatiotemporal information. Not only does this provide an opportunity to sample the feedback at multiple points of time and space, but the movement of the cursor might be attentionally engaging^15^. Third, continuous feedback is provided as soon as the movement is initiated, whereas endpoint feedback is only available when the hand reaches the target. Similar to the attenuation observed with delayed feedback^8–13^, the inherent delay in endpoint feedback with respect to movement initiation may attenuate adaptation.

The goal of the present study was to systematically investigate the difference in adaptation to continuous and endpoint feedback, assessing the relative contributions of dynamics, continuity, and onset timing. To isolate implicit adaptation, we employed task-irrelevant clamped feedback^16^. In this task, the angular divergence between the feedback and target is fixed, independent of the position (and thus movement) of the participant’s hand (Fig. 1b). Participants are fully informed of the feedback manipulation and instructed to ignore the feedback and always reach straight to the target. Despite these instructions, the participant’s behavior has all of the hallmarks of implicit adaptation: Across trials, the heading angle of the hand gradually shifts away from the target in the opposite direction of the clamped feedback, and a pronounced aftereffect is observed when the feedback is removed^16,17^. Participants are not aware of this change in behavior, believing their terminal hand position to be near the target throughout the experiment^18^.

Clamped feedback provides a unique opportunity to manipulate the timing of both continuous and endpoint feedback. Because the feedback is movement-invariant and predetermined, we can manipulate the onset, duration, and offset of the feedback. For example, endpoint feedback can be presented at movement onset, or even prior to movement onset. Moreover, to examine the influence of temporal continuity, we can manipulate the duration of the feedback to match the endpoint and continuous feedback on this dimension. Through a series of experiments, we manipulate these variables to gain insights into how sensorimotor adaptation is influenced by the spatial-temporal relationship between a movement and its associated feedback.

## Results

### Implicit adaptation is influenced by the feedback onset time

In the initial experiments, we used a web-based platform to manipulate the temporal and spatial properties of the feedback in a visuomotor rotation task (Fig. 1a)^19^. Using their trackpad, participants were instructed to make center-out ‘reaching’ movements. To elicit implicit sensorimotor adaptation, we used clamped feedback in which the cursor was always rotated from the target location by a fixed angle of 15° (Fig. 1b), and thus, not contingent on the participant’s actual movement direction.

We compared three feedback conditions in Experiment 1a (Fig. 1c). Two of these corresponded to the standard modes of feedback, continuous and endpoint, with the feedback cursor presented throughout the movement for the former and only at the endpoint for the latter. We expected to observe greater adaptation to continuous feedback compared to standard-endpoint feedback, demonstrating that this well-established effect is manifest on our web-based platform. For the third condition, we manipulated the onset time of the endpoint feedback, presenting it at the endpoint position for one refresh cycle as soon as movement initiation was detected. In this way, this early-endpoint feedback is matched to continuous feedback in terms of onset time and to the standard-endpoint feedback in terms of spatial position and temporal duration.

Following a baseline period with veridical feedback, clamped feedback was presented for 400 trials, with the three modes of feedback tested in different groups of participants. Participants showed robust adaptation in all three conditions (Fig. 2a), with the shift in hand angle persisting across a no-feedback washout block. Consistent with prior studies^4–7^, participants adapted less in response to standard-endpoint feedback compared to continuous feedback. Surprisingly, early-endpoint feedback resulted in a level of adaptation that was comparable to continuous feedback. Using a non-parametric cluster-based permutation analysis (see Methods), significant differences between conditions were observed across almost the entire extent of the perturbation and washout blocks. Focusing on a pre-specified epoch near the end of the perturbation block, we performed a series of post hoc pairwise comparisons. The hand angle in the standard-endpoint condition was lower than in the continuous (t(61)=3.2, p_bf_=0.004, BF_10_=16.3, d=0.81) and early-endpoint conditions (t(57)=3.3, p_bf_=0.004, BF_10_=19.4, d=0.85). No difference was found between the continuous and early-endpoint conditions (t(56)=0.18, p_bf_=1, BF_10_=0.27, d=0.048). This result provides a striking demonstration of the relevance of feedback onset timing: Providing endpoint feedback as early as continuous feedback, even for just a single refresh cycle, was sufficient to offset the attenuating effects of standard-endpoint feedback.

**Figure 2.**
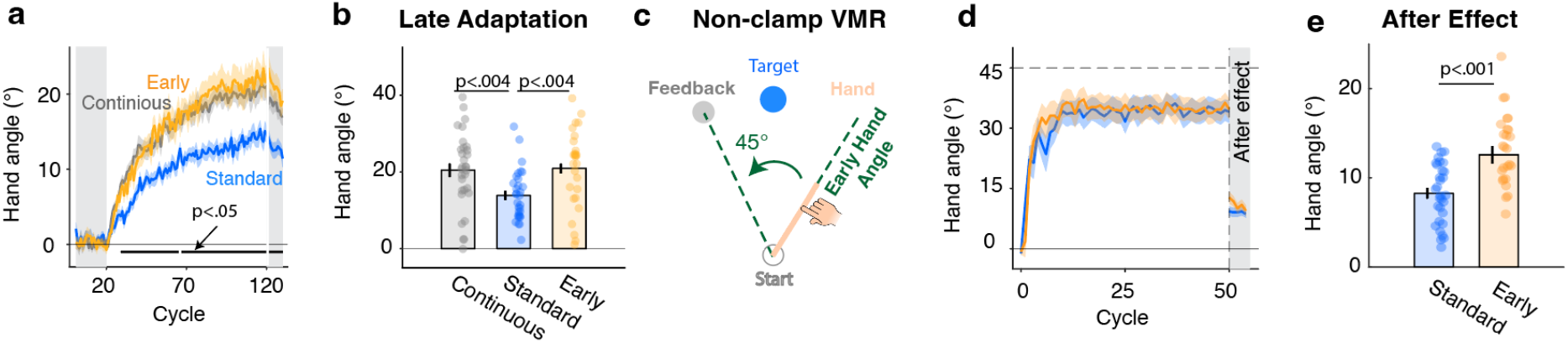
Implicit adaptation is enhanced by advancing endpoint feedback. **a**, Experiment 1a: Hand angle time course for the continuous (gray), standard-endpoint (blue), and early-endpoint (yellow) conditions in Experiment 1a. The light gray regions indicate baseline and washout (no feedback) blocks. Black horizontal lines at the bottom indicate clusters showing significant main effects of feedback. **b**, Implicit adaptation magnitude calculated over cycles 110-120 (late adaptation). **c**, Experiment 1d: Illustration of trial in visuomotor rotation task with contingent endpoint feedback; the cursor is rotated by 45° with respect to the projected position of the hand based on actual hand position early in the movement. **d**, Time courses of hand angle, and **e,** implicit adaptation magnitude calculated over the first cycle in the washout block of Experiment 1d. Shaded area in a and d and error bars in b and e represent standard error. Dots in b and e represent individual participants.

There was a near-significant difference between groups in movement time (F(2, 88)=2.9, p=0.06, BF_10_=0.49, 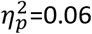) and a significant group difference in reaction time (F(2, 88)=9.4, p<0.001, BF_10_=85.9, 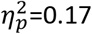). Overall, the standard-endpoint group tended to start their movements faster and move slower (Fig. S2). However, this pattern was observed in both the baseline and perturbation blocks, indicating that it was likely due to random variation between the three groups rather than due to the clamped feedback manipulation. Nonetheless, we confirmed that the advantage of continuous and early-endpoint feedback holds even when we regressed out individual differences in median movement duration and median reaction time, as well as age and gender (Table S1).

The advantage of early-endpoint feedback could occur if participants used the feedback to make mid-movement feedback corrections. For example, seeing the feedback shifted 15° in the clockwise direction might elicit an online correction in the opposite direction, a response that would inflate our estimate of adaptation. In Experiment 1a, our code only saved hand position when the target radial distance was reached, making it impossible to determine if there was evidence for online corrections. To address this concern, we recorded the entire movement trajectory for all three conditions in Experiment 1b. The basic pattern of adaptation was replicated with the early-endpoint and continuous conditions producing comparable levels of adaptation, both higher than that observed in the standard-endpoint condition (Fig. S3). We found no evidence of online corrections in all three feedback conditions, and no difference in reaction time, movement time, and movement speed (Fig. S4).

The code for the online study was written with the aim of presenting the feedback in the early-endpoint condition immediately after movement initiation. However, the actual presentation time is delayed by two factors: 1) The time to detect movement along the trackpad and 2) the time required to present the visual feedback (Fig. S1). We quantified this delay in the lab by having a few pilot participants perform the task while we made video recordings simultaneously of the participants’ hand and monitor with different devices. By this method, we estimated the delay between movement onset and feedback onset (in the early feedback condition) to be between 150-180ms. A previous study showed the delay to present visual feedback on web-based platforms across devices is around 11.5 (±15.4) ms^20^; as such, the majority of the delay in our system is likely due to a delay in detecting movement onset from a trackpad. Thus, the early endpoint feedback is likely presented mid-movement rather than at movement onset. We return to this issue in Experiment 3.

Independent of the delay in movement onset detection, the finding that early-endpoint feedback produces an adaptive response similar to that observed with continuous feedback was surprising. We wondered if this might be the result of a ceiling effect given that implicit adaptation is known to be invariant in response to a large range of errors (~10° - 90°)^16,17,21,22^. To address this question, we compared continuous and early-endpoint feedback in Experiment 1c using a small, 2° visual error clamp. As expected, the asymptote in response to this error was lower than observed in Experiment 1 where the clamp size had been 15°. Importantly, we again did not observe any difference between early-endpoint and continuous feedback (t(50)=0.93, p=0.35, BF_10_=0.40, d=0.26, Fig. S5).

In Experiment 1d we replaced the visual clamp with position-contingent feedback, addressing the concern that the boost observed with advanced endpoint feedback might be idiosyncratic to the clamp method^23^. We used a 45° rotation and compared the early-endpoint and standard-endpoint conditions. It is, of course, not possible in the early endpoint condition to precisely plot the feedback cursor at movement onset based on the (future) endpoint position of the hand. However, given that the movement trajectories are relatively straight, we could predict the endpoint position of the hand based on the heading angle sampled just after movement onset (Fig. 2c). To keep the spatial information similar across conditions, we applied the same method in the standard-endpoint condition (determined angular position of feedback based on the initial heading angle). Adaptation was larger than in experiments 1a-c for both early-endpoint and standard-endpoint conditions (Fig. 2d) as participants followed the instructions to ‘make the cursor hit the target.’ This behavior likely reflects the composite contributions of both explicit and implicit processes. To estimate implicit adaptation, we focused on the aftereffect, calculated as the mean hand angle in the first cycle of the washout block). Here we again observed greater adaptation in the early-endpoint condition compared to the standard-endpoint condition (t(56)=3.9, p_bf_<0.001, BF_10_=101.1, d=1.0, Fig. 2e).

In sum, the results of this first set of experiments demonstrate that advancing the timing of endpoint feedback produces a significant increase in implicit adaptation; indeed, the adaptive response to early endpoint feedback is comparable to that observed in response to continuous feedback. This pattern was observed with feedback signals that varied considerably in terms of error size and movement contingency (clamped or contingent). The absence of evidence of corrective movements suggests that, mechanistically, advancing the onset time of endpoint feedback enhances processes involved in adaptation of a feedforward motor plan rather than processes invoked for online corrections.

### Temporal and spatial continuity does not influence implicit adaptation

Advancing endpoint feedback ensures that feedback onset is matched between continuous and endpoint feedback conditions. However, they still differ in terms of temporal and spatial extent, with continuous feedback available for a longer duration and traversing a larger spatial distance. We next asked if these variables influence implicit adaptation. We examined the influence of temporal continuity in Experiment 2a by testing two new conditions. In one condition, we extended the presentation time of the early-endpoint feedback to match it to the duration of the entire movement. In another condition, we extended the duration of the standard endpoint to match the movement time for that trial (on average 84 ms, Fig. 3a-b). Thus, in both conditions, the static feedback is visible for the same mean duration as continuous feedback. Compared to the original conditions in Experiment 1 (1 refresh cycle), temporally extending the presentation of endpoint feedback did not influence the time course of adaptation for either the standard-endpoint or early-endpoint conditions (Fig. 3c-d). A regression model showed a significant effect of feedback onset time (coefficient 95% CI: [0.91,9.1], t(118)=2.4, p=0.017, BF_10_=9.4, 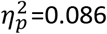) with no effect of the presentation time duration (coefficient 95% CI: [-5.3,2.8], t(118)=-0.62, p=0.53, BF_10_=0.004, 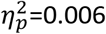).

**Figure 3.**
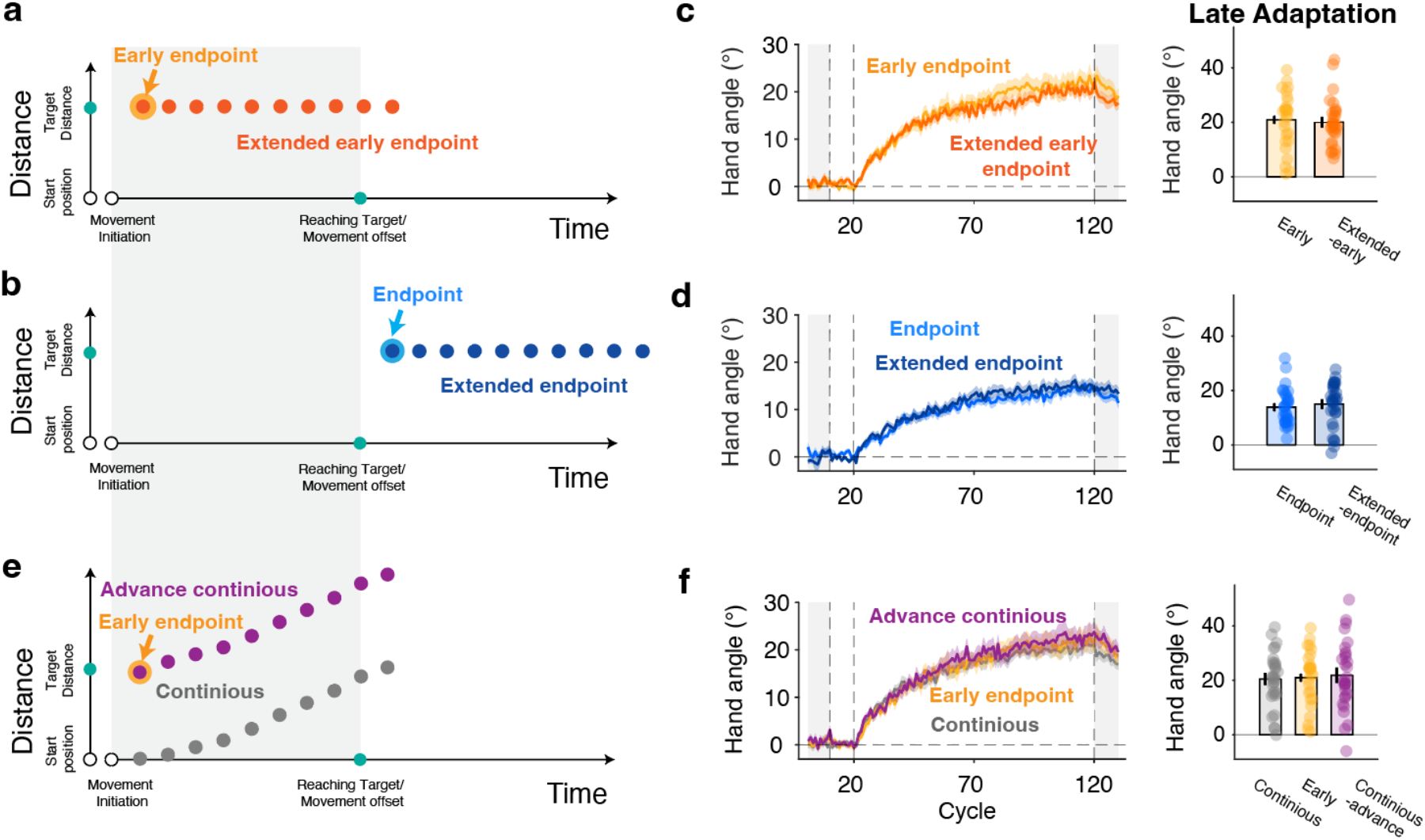
Experiment 2: Temporal and spatial extent does not influence implicit adaptation. **a-b**, Illustrations of extended versions of early- and standard-endpoint feedback conditions. X-axis indicates time and Y-axis indicates radial distance of the cursor relative to the start position. Each dot represents a refresh cycle. The gray area indicates the movement period. Feedback was presented, on average, for 5 cycles in the extended endpoint conditions. Note that the lighter colors show the timing for the single-cycle variants of early-endpoint and standard-endpoint used in Experiment 1. **c-d**, Left: Time course of hand angle in Experiment 2. The shaded area represents standard error. Light gray areas indicate baseline and washout blocks. No significant clusters were found in the comparison of the brief (1 cycle, data from Experiment 1) and extended versions. Right: Comparison of hand angle measured in late adaptation. **e**, In the advanced continuous condition, the feedback cursor appeared at the endpoint location at movement onset and then moved in the direction of the hand movement. **f**, As in c,d: No significant differences were observed in the cluster-based analysis of the learning functions or during late adaptation.

To look at the effect of spatial continuity, we created an advanced-continuous feedback condition in Experiment 2b. Here the cursor was presented at the endpoint position upon movement initiation and then moved beyond the target as the participants reached towards the target (Fig. 3e). Thus, the initial position of the feedback is matched to that of the endpoint conditions. This condition resulted in a similar extent of adaptation as standard-continuous in response to both a 15° (Fig. 3f; t(55)=0.30, p=0.77, BF_10_=0.28, d=0.079) and 2° (Fig. S5; t(42)=-1.3, p=0.21, BF_10_=0.58, d=-0.4) visual clamp, surpassing that observed with standard-endpoint feedback. These results are consistent with the hypothesis that the sensorimotor adaptation system is sensitive to the onset time of the feedback but not the temporal or spatial extent of the feedback, two fundamental features of continuous feedback.

### The optimal feedback onset time is locked to the movement onset

We next consider two hypotheses that might account for the advantage of early endpoint feedback over standard-endpoint feedback. One hypothesis centers on the idea that optimal adaptation occurs when the error signal is synchronized with the sensory prediction. Considering that 1) the motor command is generated just prior to movement onset, and 2) the sensory prediction is derived from an efference copy of the motor command^1^, we assume that the representation of the prediction is strong at a certain time after movement onset. As such, decay in this representation would be minimized when the visual feedback is advanced in time. Alternatively, implicit adaptation may be optimal when the radial position of the visual feedback is subjectively in synchrony with the position of the hand. Given that visual signals are slower than proprioceptive signals^24^, advancing the visual feedback might be a way to offset this latency difference.

To evaluate these hypotheses, we turned to a trial-by-trial design in Experiment 3. We used a 15° visual clamp with the direction of the clamp pseudo-randomized to be either clockwise or counterclockwise across trials (and thus prevent accumulated learning). With this design, the index of adaptation is the trial-by-trial change in hand angle^25^. We varied the onset time of the feedback using intervals that were designed to range from 200 ms prior to movement onset to 300 ms after movement onset. We set this window based on a running average estimate of each participant’s mean movement initiation time (Fig. 4a). To test whether the optimal feedback time is locked with the movement onset or movement offset, we manipulated movement duration by varying movement distance (7 cm or 15 cm) across participants. The median movement durations for the two conditions were 176.7ms and 272.2ms, respectively (Fig. 4b, Fig. S6). By testing the participants in the lab, all on the same apparatus, we were able to establish precise control over the timing of three critical events -- movement onset, feedback onset, and the time at which the movement reached the target amplitude.

**Figure 4.**
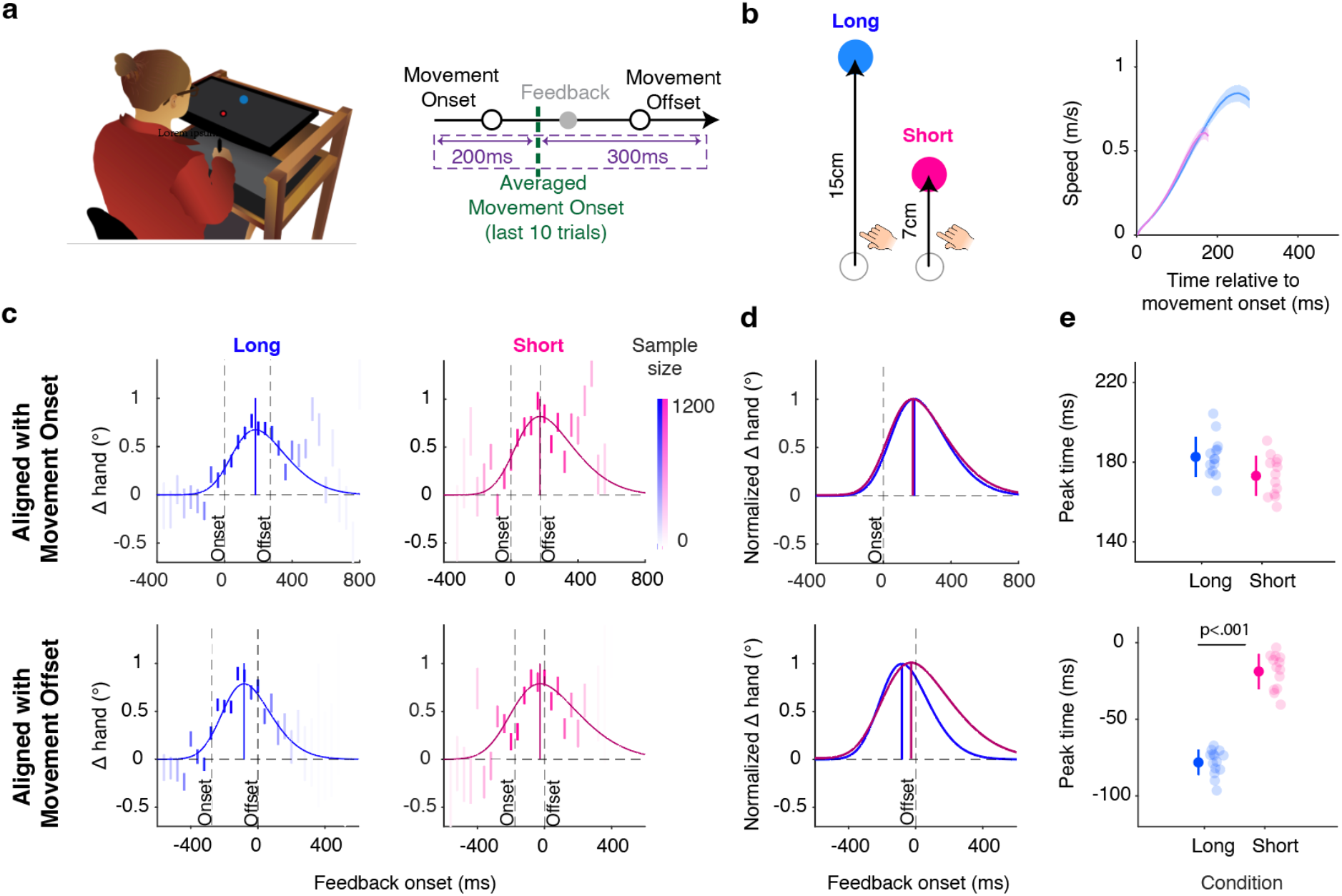
The optimal timing of endpoint feedback is associated with the movement onset. **a**. Schematic of the experimental setup (left) and procedure in the lab for Experiment 3 (right). Feedback time for each trial was predetermined, selected from a window ranging from −200 ms to +300 ms relative to the average movement onset time of the last 20 trials. Although participants were instructed to reach through the target (‘slicing’ movements), we defined ‘movement offset’ as the time when the hand reached the target distance. **b**, Left: To vary movement duration, the required movement amplitude was short or long. Right: The speed profiles for the two movement amplitudes. Shaded area represents standard error. **c**, Change of hand angle (i.e., Δ hand angle, or trial-by-trial motor correction) as a function of feedback onset time with respect to the movement onset (top row) or movement offset (bottom row) for the long and short movements. Each bar is a bin of 40 ms and the darkness of it indicates the relative number of samples in that bin. Colored curve indicates the best-fitted skewed Gaussian, with the colored vertical line marking the peak of the function. **d**, Best-fitted skewed Gaussians, normalized to peak height. **e**, Optimal feedback time relative to the movement onset (top) or offset (bottom), estimated by Jackknife resampling with each dot a sample that included 12 of the 13 participants. The optimal times are statistically indistinguishable for the short and long conditions when determined relative to movement onset, but not when determined relative to movement offset. Error bar represents the standard deviation.

Feedback onset time had a non-monotonic effect on the trial-by-trial motor correction (Fig. 4c top). Minimal adaptation is observed when the feedback leads movement onset. The function rapidly rises, reaching a peak near movement offset before falling off for longer feedback onset times. The bottom row of Figure 4c plots the same data, but now aligned to the sample at which the hand was detected to be at the target amplitude (referred to as movement offset). For the short movement, the adaptation function peaks close to movement offset. However, for the long movement, the peak is advanced, coming prior to movement offset. Strikingly, adaptation peaked at roughly the same time after movement onset for both the short and long movements.

The functions in Fig. 4c are based on group-averaged data. It is important to verify that the functions, especially the one showing that the peak occurs in advance of movement offset (e.g., long movement condition), are not distorted due to averaging movement time across trials or participants. To this end, we plotted the motor correction function as a percentage of movement duration on each trial. For example, a 50% feedback onset time corresponds to trials in which the feedback is presented at the midpoint of the movement duration, whereas 100% would indicate trials in which the feedback is presented when the hand reaches the target eccentricity. In this normalized analysis, we observed a similar pattern (Fig. S7). Importantly, the peak of the function for the long movement condition occurs in advance of when the hand reaches the target amplitude, at around 75% of the movement duration.

To quantify the peak of the feedback function, we used a model-based approach (Fig. 4c). Assuming a skewed Gaussian function, we calculated the time of peak adaptation with respect to movement onset and movement offset, comparing the functions for the short and long movement durations. With respect to movement onset, the functions for the short and long duration movements were very similar (Fig. 4d top), with peaks that were indistinguishable (long: 182.7±9.9ms, short, 173.2±10.0ms; z=0.67, p=0.50, d=0.19, Fig. 4e top). In contrast, when we compared the functions for the short and long duration movements aligned with respect to movement offset time, the peaks were markedly different (long: −78.1 ± 8.3ms; short: −18.7±11.5ms, z=4.2, p<0.001, d=1.2, Fig. 4d-e, bottom).

Taken together, those results indicate that the optimal time to present feedback does not correspond to the time at which the hand position and feedback position are aligned. Rather, the optimal time for feedback appears to be at a fixed delay relative to movement onset, consistent with the hypothesis that learning is strongest when the feedback is temporally aligned with the sensory prediction.

## Discussion

Visual feedback provides an essential source of information to improve motor performance. Continuous feedback helps the driver navigate on a curvy country road; endpoint feedback can aid a basketball player when shooting free throws. While many studies have shown that continuous feedback induces faster learning relative to endpoint feedback^4–6^, the reason for this benefit has not been clear. Not only does continuous feedback provide extended spatiotemporal information, but its onset is also earlier in the movement. In the present study, we used non-contingent, clamped feedback to examine various factors that might underlie the disadvantage of endpoint feedback. The results show that, whereas the duration and spatial extent of the feedback had no impact on the strength of adaptation, the onset time of the feedback was critical. Advanced ‘endpoint’ feedback, even when limited to a single frame, resulting in adaptation comparable to that observed with continuous feedback.

These observations are especially surprising given that continuous feedback provides richer information than endpoint feedback. However, various lines of studies indicate that implicit adaptation is relatively insensitive to the quality of the feedback. For example, adaptation is insensitive to the uncertainty of the visual feedback, at least for relatively large errors^22^, suggesting that the implicit learning system may not be sensitive to the quality or ecological validity of the feedback. This point is further underscored by the very fact that robust adaptation is observed in response to clamped feedback despite participants’ awareness of the manipulation. These observations point to a system that is highly modular, automatically using an error signal to make feedforward adjustments to keep the sensorimotor system precisely calibrated. The benefit of continuous feedback may be pronounced in feedback control, enabling online adjustments to ensure that the movement goal is achieved^26,27^.

The importance of feedback timing has been highlighted in prior studies of visuomotor adaptation. That body of work has emphasized how delaying endpoint feedback can dramatically attenuate adaptation^8,9,11,13,28^. Implicit in this work is the assumption that the optimal time for endpoint feedback is at movement offset, that is, when the feedback is temporally and spatially synchronized with the hand ^13,29^. Because these studies used position-contingent feedback, it was only possible to delay the feedback. Taking advantage of the fact that position-independent, clamped feedback is effective in eliciting adaptation, we were able to temporally advance ‘endpoint’ feedback. The enhancement of learning observed with this method indicates that the advantage of continuous feedback does not rest on its spatiotemporal continuity, and that it is not essential that the position of the feedback be synchronized with the position of the hand.

Having demonstrated the advantage of early endpoint feedback, we set out to determine the optimal time for the feedback. The clamp method allowed us to parametrically manipulate the onset time of endpoint feedback. Here, we transitioned from a web-based platform to the lab to minimize measurement delays and test a wider range of values ranging from well before the movement onset to beyond movement offset. Moreover, by using two distinct movement durations (by manipulating movement amplitude), we could examine if optimal timing was linked to movement onset or movement offset. We observed non-monotonic functions for both amplitudes. The attenuation for the longest feedback onset latencies provides another demonstration of the cost of delayed feedback. Moreover, the attenuation for the shortest latencies indicates that there is a cost for presenting the feedback too early, including the time of movement initiation.

The peak of the function (i.e., the optimal timing for feedback) was observed at a midway point, one in which the position of the feedback was in advance of the position of the hand. One hypothesis to account for this effect is based on models of multisensory integration. In this framework, the perceived hand position, the signal essential for computing the error, is an integrated representation based on a variety of inputs, including vision and proprioception^30^. Temporally, one would expect that the contribution of the visual signal will be strongest when it is synchronized with the proprioceptive signal. In terms of neural responses in the brain, visual inputs are delayed by approximately 50 ms relative to proprioceptive inputs^24,31,32^. Advancing endpoint visual feedback by this interval could enhance visual-proprioceptive synchronization, and thus boost learning. Importantly, since movement offset is defined by hand position, this hypothesis predicts that the optimal timing of feedback should be constant with respect to movement offset. However, this prediction was not supported by the results. More generally, the brain has likely evolved mechanisms to compensate for inherent differences in transmission delays across sensory modalities, negating the need for temporal synchronization in deriving a multisensory signal ^33^.

Whereas the multisensory integration hypothesis focuses on the observed feedback, an alternative hypothesis focuses on how this information is compared to the predicted sensory outcome. The latter is assumed to be generated from an efference copy of the motor command. We hypothesize that the feedback timing function reflects the strength of the representation of the sensory prediction. When considered as a discrete event, the 170 ms delay may reflect the interval between the efference copy and time in which the prediction is available; when considered as a continuous neural process, the representation of the prediction may reach its maximal strength around 170 ms after movement initiation. By either view, we assume this representation will decay after its peak. When feedback is temporally advanced, adaptation is therefore strengthened since the feedback arrives prior to the decay of the sensory prediction. This hypothesis is consistent with the observation that the optimal time was time-locked to movement onset, independent of movement duration.

Temporal constraints are a prominent feature of cerebellar-dependent learning, including sensorimotor adaptation and eyeblink conditioning^33^. In the latter, the animal learns to make a blink in response to an arbitrary stimulus (conditioned stimulus, CS) that is predictive of an aversive event (unconditioned stimulus, US). This form of learning is highly sensitive to the interval between the CS and the US^34^, showing a non-monotonic function similar to that observed in the present study. Learning is negligible when the US occurs before or together with the onset of the CS, peaks when the CS leads the US by between 200 – 400 ms, and decreases for longer intervals^35–38^. The rise of this function has been assumed to reflect the time required to generate an expectancy of the US and adaptive motor response that will attenuate the aversive effects of the US. The reduced efficacy of learning for longer CS-US intervals is assumed to reflect temporal limitations within the cerebellum for maintaining the sensory prediction. This account of the optimal timing for eyeblink conditioning is similar to our account of optimal timing for visuomotor adaptation. In the latter, the motor commands and visual feedback serve as equivalents for the CS and US, respectively. Consistent with this notion, we have recently shown that when temporal constraints are imposed, two signature phenomena of classical conditioning, differential conditioning and compound conditioning, are observed in visuomotor adaptation^39^.

It remains to be seen whether the benefits of advanced timing hold over a broad range of contexts. Our experimental manipulations were limited to relatively simple reaching movements, performed over two movement amplitudes. Future behavioral studies should examine the effect of feedback timing for movements that span a wider range of durations and contexts. Indeed, a recent study suggests that our optimal timing hypothesis could be tested in the absence of movement. By using a Go-NoGo task, Kim et al. observed adaptation in response to clamped feedback even after trials in which the movement was aborted^40^. Presumably, a motor command was generated on the no-go trials, resulting in a sensory prediction that could be compared to the clamped feedback. This task might be ideal for examining feedback timing given that movement kinematics are eliminated on the no-go trials, removing additional and potentially conflicting sources of information (e.g., proprioceptive signals from the moving limb).

### Conclusions

A core principle featured in motor learning textbooks is that endpoint feedback elicits less learning than continuous feedback^41,42^. The present results indicate that a major reason for the disadvantage of endpoint feedback is that it becomes available later than continuous feedback; when ‘endpoint’ feedback is temporally advanced, implicit adaptation was enhanced, reaching a level comparable to that observed with continuous feedback. By systematically varying the onset time of ‘endpoint’ feedback, we found that the optimal feedback time was time-locked to movement onset rather than movement offset. We hypothesize that adaptation is optimized when the sensory prediction is at maximal strength for comparison with the sensory feedback in generating an error signal. These results underscore novel temporal constraints underlying cerebellar-dependent sensorimotor learning.

## Methods

### Participants

Testing was conducted online for Experiments 1-2 and in the lab for Experiment 3. For online studies, a total of 272 participants (116 female, mean age = 24.5, SD = 4.7) were recruited through the website prolific.co. We recruited 34 participants for each condition based on a power analysis (see Supplementary Methods). After eliminating participants who failed to meet our performance criteria (see below), the analyses were based on data from 239 participants (27-32 for each condition, 93 females, mean age = 24.3, SD = 4.5). Based on self-report data from a prescreening questionnaire, all of the participants were right-handed with normal or corrected-to-normal vision. The online participants were paid around $8/h. For the lab-based experiment, we recruited 26 undergraduate students (15 female, mean age = 21.5, SD = 4.5) from the University of California, Berkeley community. All of the participants were right-handed based on their scores on the Edinburgh handedness test^43^ and had normal or corrected-to-normal vision. These participants were paid $15/h. All experimental protocols were approved by the Institutional Review Board at the University of California, Berkeley. Informed consent was obtained from all participants.

### Web-based experiments

Online experiments (Exp 1-2) were performed using our web-based experimental platform, OnPoint^19^. The code was written in JavaScript and presented via Google Chrome, designed to run on any laptop computer. Visual stimuli were presented on the laptop monitor and movements were produced on the trackpad. Data were collected and stored using Google Firebase. The experimental session lasted around 40 minutes

### Procedure

#### Experiment 1a

Clamp rotation task. To start each trial, the participant moved the cursor to a white start circle (radius: 1% of the screen height) positioned in the center of the screen. After 500 ms, a blue target circle (radius: 1% of the screen height) appeared with the radial distance set to 40% of the screen size. There were four possible target locations (+/-45°, +/-135°). The participant was instructed to produce a rapid, out-and-back movement, attempting to intersect the target. The target disappeared when the amplitude of the cursor movement reached the target distance. To help guide the participant back to the start location, a white cursor (radius: 0.6% of screen height) appeared when the hand was within 40% of the target distance. If the movement time was >500 ms, the message ‘Too Slow’ was presented on the screen for 500ms.

We used a visual clamp to elicit implicit sensorimotor adaptation, manipulating feedback onset time, presentation duration, and spatial continuity. Three types of clamp feedback were employed in Experiment 1 (between-subjects). (1) Continuous feedback: The radial location of the cursor was based on the radial extent of the participant’s hand and was visible during the whole movement (up to reaching the target distance) but was independent of the angular position of the hand. (2) Standard endpoint feedback: The cursor was presented at the target distance for one refresh cycle (10~20ms depending on the monitor refresh rate, 16.7ms for 60 Hz monitor) after the hand reached the target distance. In this manner, the timing and position of the cursor was the same in this condition as the last frame for the continuous feedback condition. (3) Early-endpoint feedback: The cursor fleshed at the target distance for one cycle when the hand was detected to exit the start circle. Thus, the onset of the feedback cursor is at the same time as the onset time for continuous feedback.

There was a total of 520 trials in Experiment 1, arranged in four blocks. 1) A no-feedback baseline block (40 trials). 2) A feedback baseline block with veridical continuous feedback (40 trials). 3) A learning block with clamped feedback (400 trials), where the cursor followed a trajectory that was displaced at a fixed angle from the target. Right before the learning block, a set of instructions was presented to describe the clamped feedback. The angle was set to 15° and the direction of the clamp was either clockwise (CW) or counterclockwise (CCW) with respect to the target, counterbalanced across participants. The participant was informed that the cursor would no longer be linked to their movement, but rather would follow a fixed path on all trials. The participant was instructed to always reach directly to the target, ignoring the cursor. To make sure the participant understood the nature of the error clamp, the instructions were repeated. Moreover, after the first 40 trials with clamped feedback, an instruction screen appeared asking the participant to indicate if they were aiming for the target or another location. If the participant indicated they were reaching to another location, the experiment was terminated. 4) A no-feedback washout block (40 trials). Within each block, trials were grouped into cycles of four trials, with each target (+/-45°, +/-135°) appearing once per cycle (order randomized across cycles).

#### Experiment 1b-c

The methods for Experiments 1b and 1c were essentially the same as in Experiment 1a. The only change in Experiment 1b is that the whole movement trajectory was recorded. In Experiment 1c, the clamp size was reduced to 2° and we replaced the standard-endpoint condition with advanced-continuous feedback condition (see Experiment 2 below).

#### Experiment 1d

##### Visuomotor rotation task (VMR)

To confirm that the results obtained in Experiments 1a-c were not idiosyncratic to clamped feedback, we used a standard visuomotor rotation task in Experiment 1d. Here the position of the feedback cursor during the adaptation block was contingent on the participant’s hand position. To encourage strategy use, we opted to use a large 45° rotation (CW and CCW counterbalanced across participants), limited the target position to two locations (135°/315°), and instructed participants to ‘make the cursor hit the target’ ^44^.

For both endpoint and early-endpoint feedback, the position of the feedback was rotated 45° from the hand angle obtained at the second data point collected after movement initiation. Early-endpoint feedback was presented right after this data point was sampled, while endpoint feedback was presented when the radial position of the hand reached the target distance.

As in Experiment 1a, there were four blocks: No-feedback baseline (20 trials), feedback baseline (40 trials), adaptation with contingent rotated feedback (200 trials), and no-feedback washout (20 trials). Within each block, the trials were grouped into cycles of four trials, with each of the two target positions presented twice per cycle. Prior to the start of the adaptation block, the instructions described the size and direction of the rotation and emphasized that the participant should adjust their aim to compensate for the perturbation and make the cursor hit the target. Prior to the washout block, the participant was instructed to cease using any aiming strategy and to again reach directly to the target.

#### Experiment 2

We tested three additional feedback conditions in Experiment 2 to examine the effect of dynamics and feedback duration. In Experiment 2a we examined the role of feedback duration, extending the duration of the static feedback to approximate that observed with continuous feedback. In the extended-endpoint condition, the cursor appeared after the hand reached the target distance and remained visible for an interval equal to the movement duration for that trial. In the extended-early-endpoint condition, the cursor appeared at movement onset and remained visible until the hand reached the target distance. In Experiment 2b we examined the effect of advancing continuous feedback. The cursor appeared at the endpoint position as soon as movement initiation was detected and then continued along that ray until the hand reached the target distance. Thus, the position of the cursor was advanced relative to the position of the hand. The procedure was the same as in Experiment 1a and we compared the performance of these three groups to the relevant conditions from Experiment 1a.

#### Experiment 3

Experiment 3 was designed to derive a function describing how feedback timing modulates the strength of adaptation. To ensure the precise timing of the trial events, we conducted this experiment in person, using the same apparatus for each participant. Participants performed a center-out reaching task on a digitizing tablet (Wacom Co., Kazo, Japan) which recorded the motion of a digitizing pen held in the hand. Stimuli were displayed on a 120 Hz, 17-inches monitor (Planar Systems, Hillsboro, OR) that was mounted horizontally above the tablet, obscuring vision of the arm. The experiment was controlled by a Dell OptiPlex 7040 computer (Dell, Round Rock, TX) running on a Windows 7 operating system (Microsoft Co., Redmond, WA) with custom software coded in MATLAB (The MathWorks, Natick, MA) using Psychtoolbox extensions.

The start position (radius: 4 mm) was located in the lower quarter of the screen at the midline. To allow the trial-by-trial analysis of adaptation (see below), we used a single target location (radius: 7 mm, fixed at 45°). The radius from the start position to the target location was set to 7 cm for half of the participants and 15 cm for the other half of the participants. This manipulation was included to produce different movement times for the two groups of participants. During the inbound portion of the movement, a white circle was visible at the start position with the radius of the circle indicating the participants’ distance from the start position. In this way, participants were guided to the start position without directional feedback of their movement.

The experiment began with a block of 16 trials in which a cursor (radius: 3 mm) provided continuous feedback. This was followed by an extended block of 1200 trials with clamped feedback. The clamp was offset from the target by 15°, with CW and CWW deviations intermixed within a cycle of 4 trials. By mixing CW and CWW clamps, there is no cumulative effect of adaptation; rather, the dependent variable of adaptation was the trial-by-trial change in hand angle^25^. Prior to the onset of the clamped feedback block, participants were fully informed of the clamp manipulation and instructed to always move directly towards the target. The onset time of the clamped feedback was randomly sampled from a uniform distribution ranging from −200 to 300 ms relative to a running average of the individual’s movement onset time, calculated over the last 20 trials. The clamp was presented as endpoint feedback for two refresh frames (approximately 16 ms). Note that we did not impose any constraint on movement onset time.

### Data analysis

The initial data analyses were conducted in MATLAB 2020b. Hand angle was calculated as the angular difference between the target and the hand position at the target radius. Positive values indicate hand angles in the opposite direction of the perturbation experienced by that participant, the direction one would expect due to adaptation. For Experiments 1-2, movement initiation is defined as the first sample where the hand surpassed the radius of the start position. For Experiment 3, movement initiation was defined as the first sample in the time series in which sample-to-sample acceleration remained positive for displacements greater than 5 mm^45^. Movement offset was defined as the first sample in the time series in which the radial distance of the hand reached the target distance. Note that we defined movement offset as the time when the radial distance of the movement reached the target distance even though the actual end of the movement was beyond this point (i.e., slicing movement). Trials with a movement duration longer than 500 ms or an error larger than 70° were excluded from the analyses. We excluded the entire data from participants who had less than 70% valid trials (see Supplementary Methods for details).

For Experiments 1a-c and 2, the data were averaged over cycles (4 trials/cycle). We examined learning at two time points, during the late phase of the adaptation block and during the aftereffect block. Late adaptation was defined as the mean hand angle over the last 10 cycles of the perturbation block, minus the mean of the no-feedback baseline block (to adjust for individual biases in reach direction). Aftereffect was defined as the mean hand angle over the washout block, minus the mean of the no-feedback baseline block. As a continuous measure of adaptation, we used a cluster-based permutation test (Arnal et al., 2015; Fell et al., 2011), a method traditionally used to analyze the data with temporal dependencies such as EEG, and has recently been applied to learning functions ^46,47^. In Experiment 1d, late learning is contaminated by the contribution of aiming strategies. As such, we focused on the aftereffect data, comparing the heading angle in the first cycle of this block with the average heading angle during the no-feedback baseline block.

In Experiment 3, adaptation was defined as the difference in hand angle between trial *n+1* and trial *n.* To construct functions describing how adaptation changed as a function of feedback timing we computed, for each trial, the actual interval between feedback onset time and either movement onset or movement offset. For the movement onset (offset) function, negative and positive values indicate that the feedback preceded or followed movement onset (offset), respectively.

To quantify the peak in the four adaptation functions (two distances, with one function for movement onset and one for movement offset), we combined the data across trials and participants and fit the data with a skewed-Gaussian function:

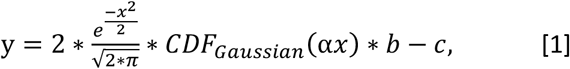

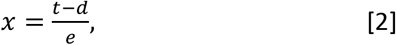

where *y* is mean Δ hand angle, *t* is the feedback onset time subtracted by either movement onset or movement offset, and *cdf_Gaussian_* is the cumulative distribution of the standard Gaussian distribution. There are five free parameters: *a, b, c, d*, and *e*, corresponding to the width, height, lower boundary, shift of the mean from zero, and skewedness of the function. To estimate the variability of each parameter, we performed a jackknife resampling procedure, leaving out the data from one participant and using the remaining data to estimate the function. This procedure was repeated with each participant excluded once.

To determine the movement trajectory, the radial axis was evenly divided into 150 segments from the initial hand position to the target. We used interpolation to obtain the heading angle for each segment. For Experiment 2, the initial heading angle was calculated by averaging the theta-angular-value at the first 30 cut points, and the end angle is defined as the theta-value at the 150^th^ cut-point (target distance).

Between-condition comparisons were performed with t-tests or ANOVAs, with Bonferroni corrections for multiple comparisons applied when appropriate. For the t-tests, we report the Bayes factor, reflective of the ratio of the likelihood of the alternative hypothesis (H1) over the null hypothesis (H0), and Cohen’s d, a measure of effect size. For ANOVAs, the effect size is reported using partial eta squared 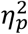. In all tests, we confirmed that the data met the assumptions of a Gaussian distribution and homoscedasticity.

## Data and code availability

Data for this paper and codes for analyses are available at https://osf.io/ej4ba/

## Author contributions

T.W., G.A., J.S.T., R.B.I. contributed to the conceptual development of this project. T.W. and T.T. collected the data. T.W. analyzed the data, prepared the figures, and wrote the initial draft of the paper, with all the authors involved in the editing process.

## Supplementary Material

### Supplementary Methods

#### Power analysis

We are interested in what is the key factor that caused the difference in implicit adaptation between the endpoint and the continuous feedback. For the block design experiments (Exp 1-2), we computed minimum sample sizes based on the no-feedback washout block from Taylor, Krakauer, Ivry (2014) in a study that used endpoint and continuous feedback. We estimated the power for an independent samples t-test using a two-tailed test with significance set at 0.05 and a power of 0.9. The effect size was d=0.91 (continuous feedback group had a mean of 25.9° S.D. of 4.9°; endpoint feedback group had a mean of 21.6° and S.D. of 4.1°), indicating a minimum sample size of 27 participants for each condition. Given our experience that some participants perform poorly (e.g., fail to pay attention) on web-based experiment (see details in the next section), we decided to recruit 25% more participants than the size suggested by the power analyses, resulting in a target of 34 participants for each feedback condition. Half experienced a clockwise perturbation and the other half a counter-clockwise perturbation. For the 2° clamp task (Exp. 1c), we recruited 28 participants for each condition with 14 people in each perturbation direction. For Experiment 3, we used a sample size (12 for each condition) that is typical in sensorimotor learning experiments.

#### Outlier removal

To minimize online corrections, we instructed the participant to move quickly. We excluded trails with a movement time longer than 500 ms. We also excluded trials in which the hand angle at the end of the movement was more than 70° from the target, under the assumption that the participant moved to the wrong target on these trials. If the total number of excluded trials was greater than 30% (movement time and direction), the entire data set for that participant was not included in the analyses. These participants either ignored the instruction to move fast or tended to repeatedly move to the same location, independent of the target location.

**Table S1.** Summary of participants’ information on web-based experiments.

|  | <u>Exp1b</u> |  |  | <u>Exp1d</u> |  | <u>Exp2a</u> |  | <u>Exp2b</u> |
| --- | --- | --- | --- | --- | --- | --- | --- | --- |
|  | Continuous | Standard-Endpoint | Early-endpoint | Endpoint | Early-endpoint | Extend endpoint | Extend early-endpoint | Advance continuous |
| Participants meeting inclusion criteria | 31 | 31 | 28 | 30 | 27 | 32 | 30 | 30 |
| Female | 19 | 9 | 7 | 13 | 11 | 14 | 13 | 17 |
| Age (mean (SD)) | 24.8(5.4) | 24.4(5.1) | 23.0(4.7) | 25.9(4.7) | 23.8(4.4) | 23.0(3.5) | 25.1(4.8) | 24.2(4.5) |
| CCW | 14 | 16 | 14 | 17 | 14 | 15 | 17 | 15 |

|  | <u>Exp1b</u> |  |  | <u>Exp1c</u> |  |  |
| --- | --- | --- | --- | --- | --- | --- |
|  | Continuous | Standard-Endpoint | Early-endpoint | Continuous | Early-endpoint | Advance continuous |
| Participants meeting inclusion criteria | 28 | 34 | 31 | 27 | 25 | 24 |
| Female | 12 | 22 | 6 | 18 | 14 | 14 |
| Age (mean (SD)) | 25.3(5.1) | 24.8(5.7) | 25.3(4.5) | 24.1(4.0) | 26.3(5.3) | 29.3(8.3) |
| CCW | 13 | 17 | 15 | 13 | 13 | 14 |

### Supplementary Results

There are substantial delays with a web-based experimental system. One delay is introduced by the time required to detect movement along a trackpad, essential for determining movement onset. A second delay is introduced by the time required to present the stimulus (feedback) on the monitor after the command is made by the code. To quantify these delays, we used a camera (frame rate = 60 Hz) to simultaneously record the monitor and movement when participants performed the web-based experiment. We performed these recordings on different devices and, for each setup, measured from the video recordings the delay between movement onset time and feedback onset time (with the program set to present feedback as soon as movement onset was detected, the early-endpoint condition). On average, the delay was 166.7 ms (Fig. S1). Note that in our setup, we cannot partition this value into that associated with movement onset detection and presentation delay. However, a previous study showed that, across devices, the delay to present visual feedback on web-based platforms is around 11.5 (±15.4) ms ^20^; as such, the majority of the delay in our system is likely due to a delay in detecting movement onset from a trackpad.

A 166.7 ms system delay in the web-based experiments would suggest that what we refer to as ‘early-endpoint feedback’ is actually occurring well into the movement. Indeed, this value is close to the peak of the motor correction function measured in the lab-based experiment (Exp. 3). Since movement ‘offset’, defined as the time when hand reached the target distance, is in the middle of the movement, the time at which movement offset is detected is not influenced by the delay in detecting movement onset. Assuming that the presentation delay for the early-endpoint feedback and the standard-endpoint feedback is the same, standard-endpoint feedback was presented around 94 ms (movement duration) after early-endpoint feedback, or around 260 ms (94+166ms) after movement onset. Therefore, the results showing early-endpoint feedback induced greater adaptation compared to standard-endpoint feedback is consistent with the motor correction function measured in the lab-based experiments.

There is also a delay between the time when the computer issues a command (e.g., ‘draw endpoint feedback) and the time at which the stimulus appears on the screen. For the lab-based experiment, Psychotoolbox adjust this for this when providing information on when the stimulus appears on the monitor. As such, we assume system delays are negligible in Experiment 3.

**Figure S1.**
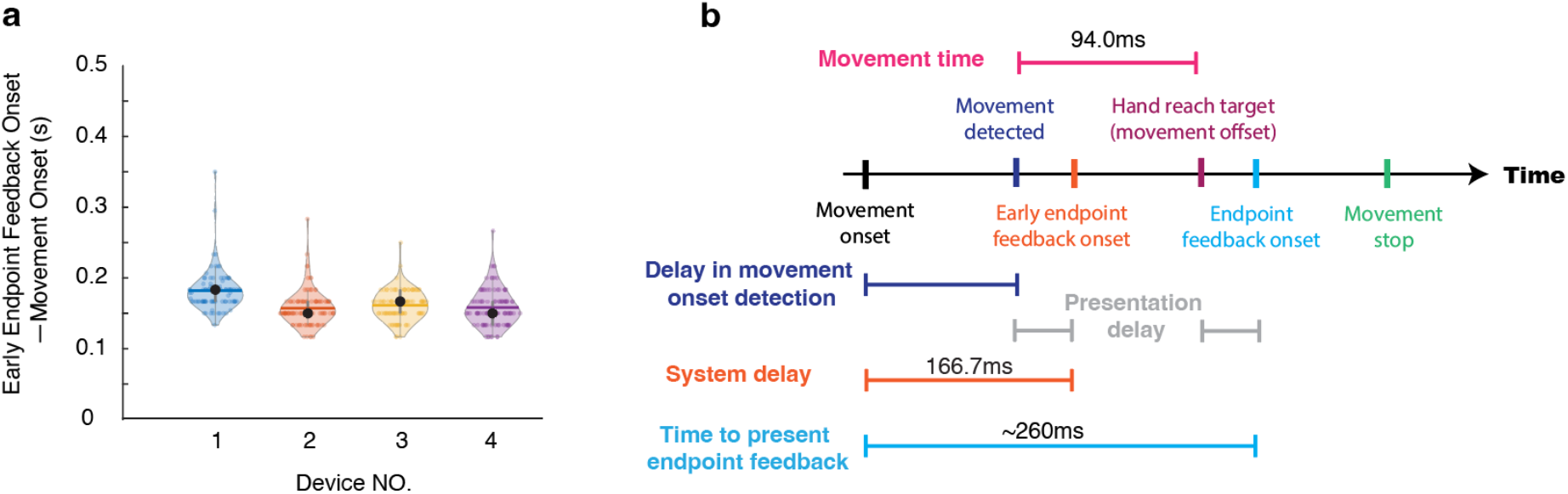
Delays in our web-based experimental system. **a,** Average system delays based on video measurements obtained when running web-based protocol on four devices (1, ThinkPad X1, 2021; 2, hp pavilion 15z-eh000; 3, MacBook Pro 2020; 4, Dell Xps 13 9365). Videos were examined and manually marked to identify the first frame in which the hand moved and the first frame in which the feedback cursor was detected. We measured 100 trials for each device. Each colored dot represents a trial. Black dots indicate the group median and the horizontal lines indicate the group means (182.2, 157.2, 161.3, 158.2 ms for devices 1-4 respectively). **b,** Timeline of the web-based experiment. The early-endpoint feedback is presented approximately 166.7 ms after movement onset. This value includes delays associated with movement onset detection and feedback presentation. The temporal difference between the standard endpoint feedback and the early endpoint feedback should be equal to movement duration. From this, we can infer that endpoint feedback is presented, on average, 260 ms after movement onset.

**Figure S2.**
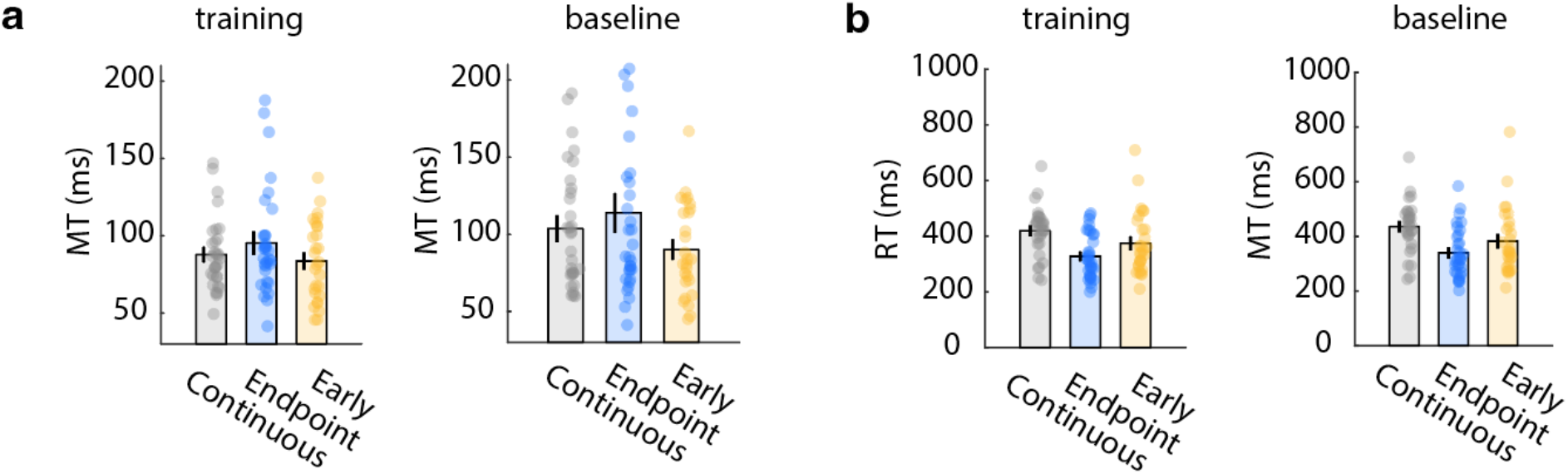
Movement duration (a) and reaction time (b) for Experiment 1. The endpoint group had longer movement times and faster reaction times compared with the other two groups. These differences are observed in both the adaptation and baseline blocks. Given that the task was identical for all three groups in the baseline block, the differences here presumably reflect random variation in our three samples and do not result from the feedback manipulation. Note that, given the delay in detecting movement onset, the values reported here overestimate reaction time and underestimate movement time.

**Figure S3.**
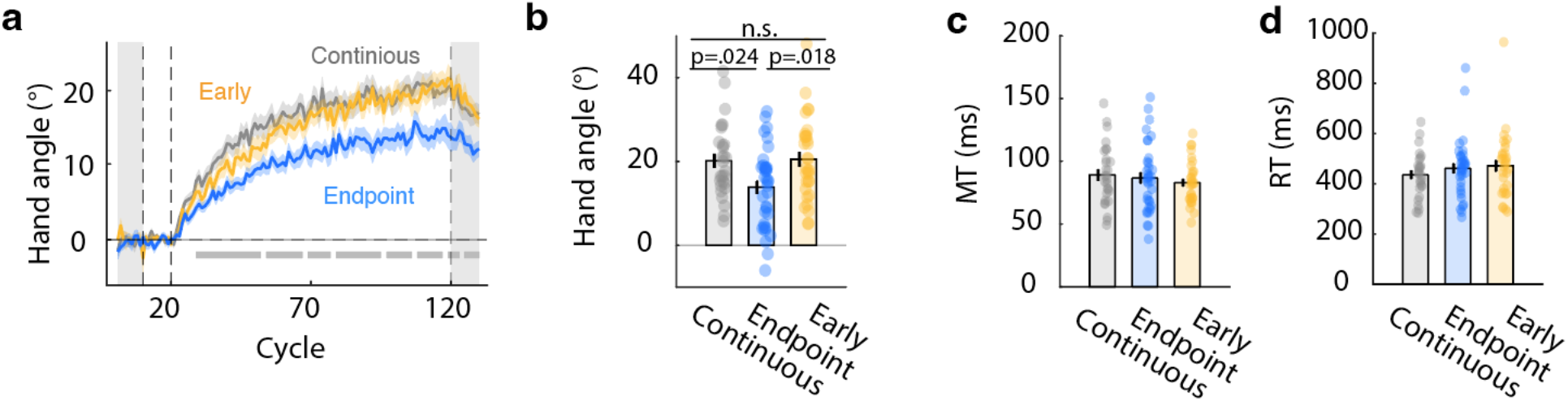
Results from Experiment 1b. Replication of Experiment 1 that includes measurement of the full trajectory. (a) Time course of hand angle. The shaded area represents standard error. The light gray areas indicate the baseline and washout blocks. Horizontal grey lines at the button indicate results of cluster-based ANOVA. Similar to the results of Experiment 1a, early-endpoint feedback induced larger adaptation compared to standard-endpoint feedback. No difference was observed between the early-endpoint and continuous feedback conditions. (**b**) Late adaptation. Error bars indicate standard error and dots represent each participant. n.s., non-significant. (**c**) Movement time is comparable across conditions (86.2±23.1ms; F(2, 90)=0.68, p=0.51, BF_10_=0.07, 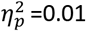). (**d**) Reaction time is comparable across conditions (457.2 ± 114.8ms; F(2, 90)=0.71, p=0.49, BF_10_=0.07, 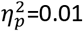).

**Figure S4.**
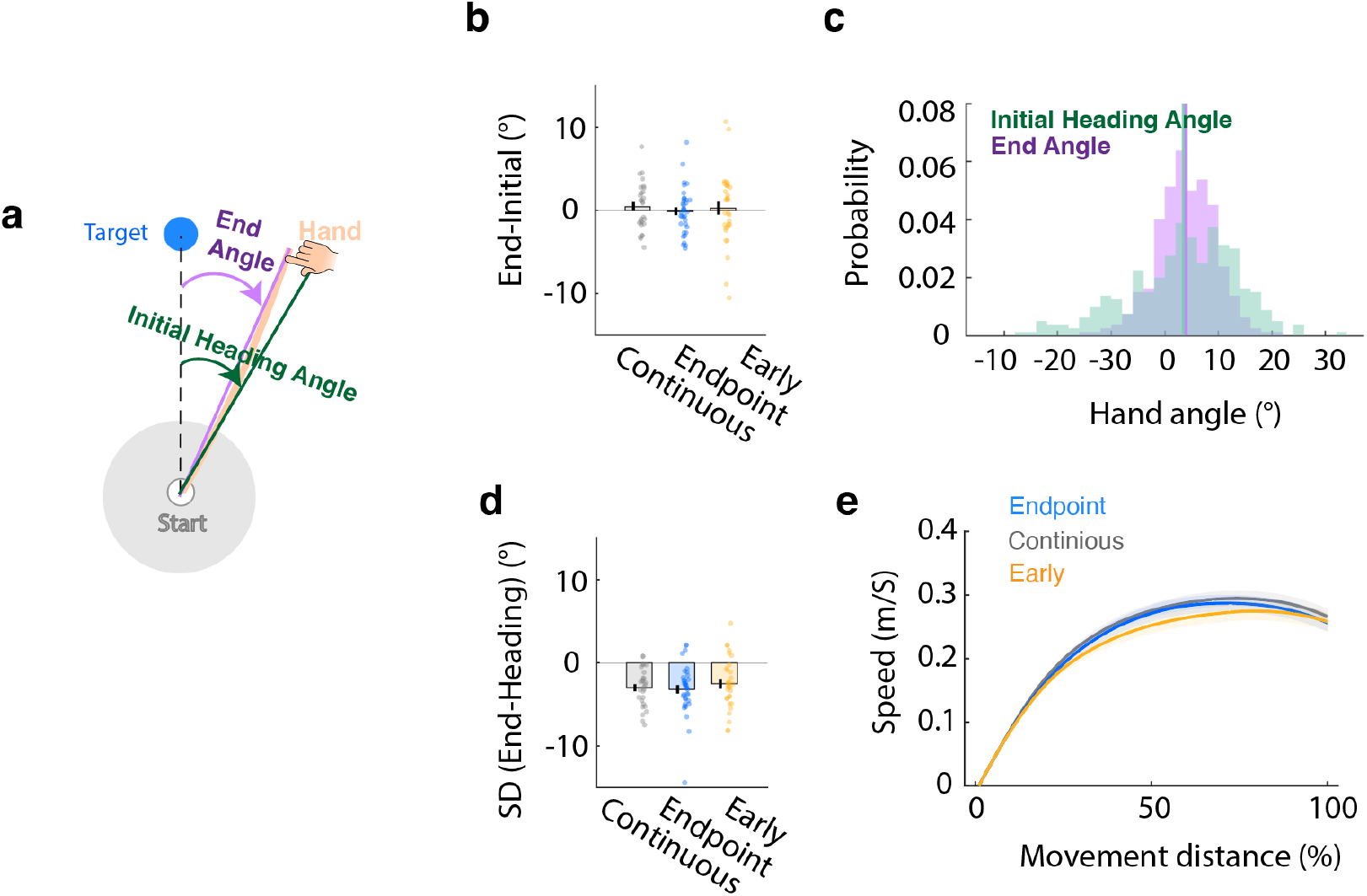
Movement kinematics were not influenced by feedback format. (a) Illustration of the definition of initial heading angle and end angle of the movement. (**b**) Change between initial heading angle and final heading angle. Positive values indicate angle of hand at target amplitude is larger than initial measurement of hand angle, the direction expected if participants were making an on-line correction. The heading angle remained relatively constant across the movement in all three conditions. Error bar indicates standard error and each dot the value for a participant. (**c**) Distribution of the initial and end heading angles for a typical participant. The initial heading angle is nosier than the end angle. (**d**) Difference in standard deviation of the initial and end heading angles. The latter is smaller in all three conditions. This effect is likely due to variation in the exact position of the hand within the start circle. Given the difference in variance and absence of evidence of online corrections, we opted to use the end angle as the primary dependent variable. (**e**) Normalized mean speed profile for the three feedback conditions.

**Figure S5.**
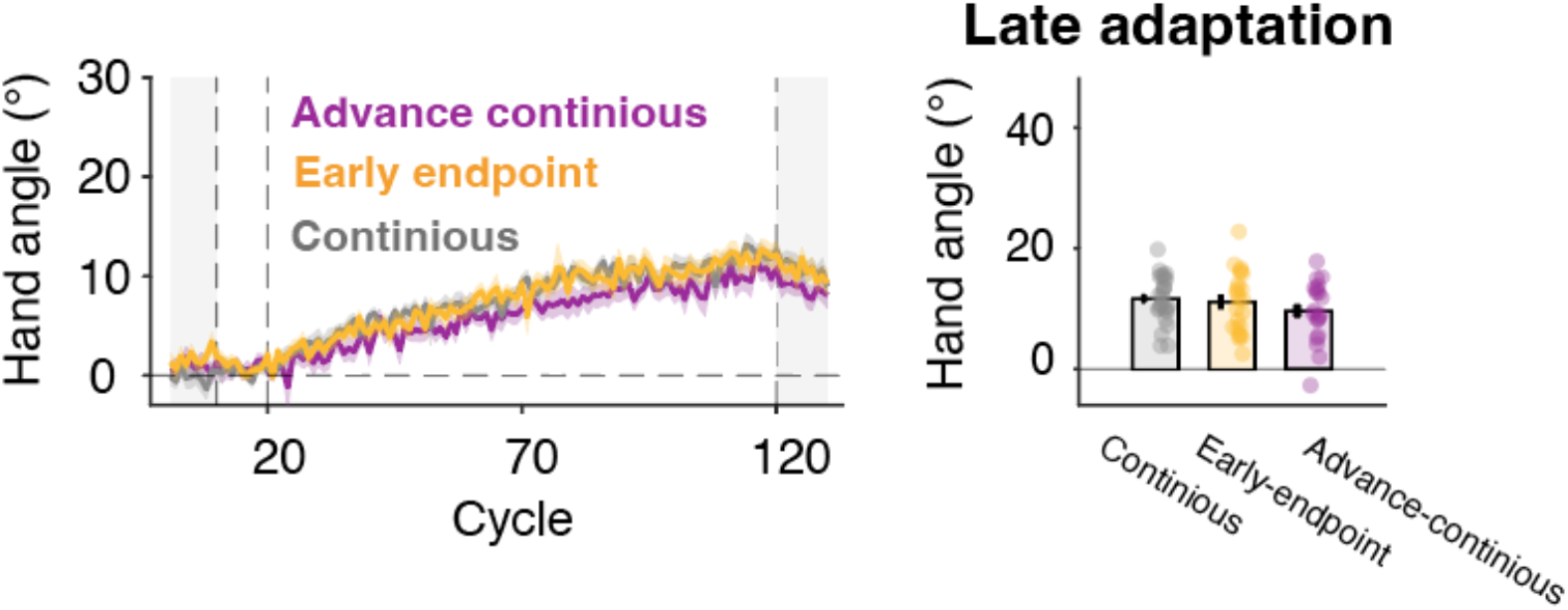
Results from Experiment 1c in which the clamp size was reduced to 2°. Adaptation was similar in response to early-endpoint, continuous, and advanced-continuous feedback conditions. Left: Time course of hand angle. The shaded area indicates standard error. No significant differences were found in the cluster-based ANOVA. Right: Hand angle in late adaptation. Error bar indicates standard error and each dot a participant.

**Figure S6.**
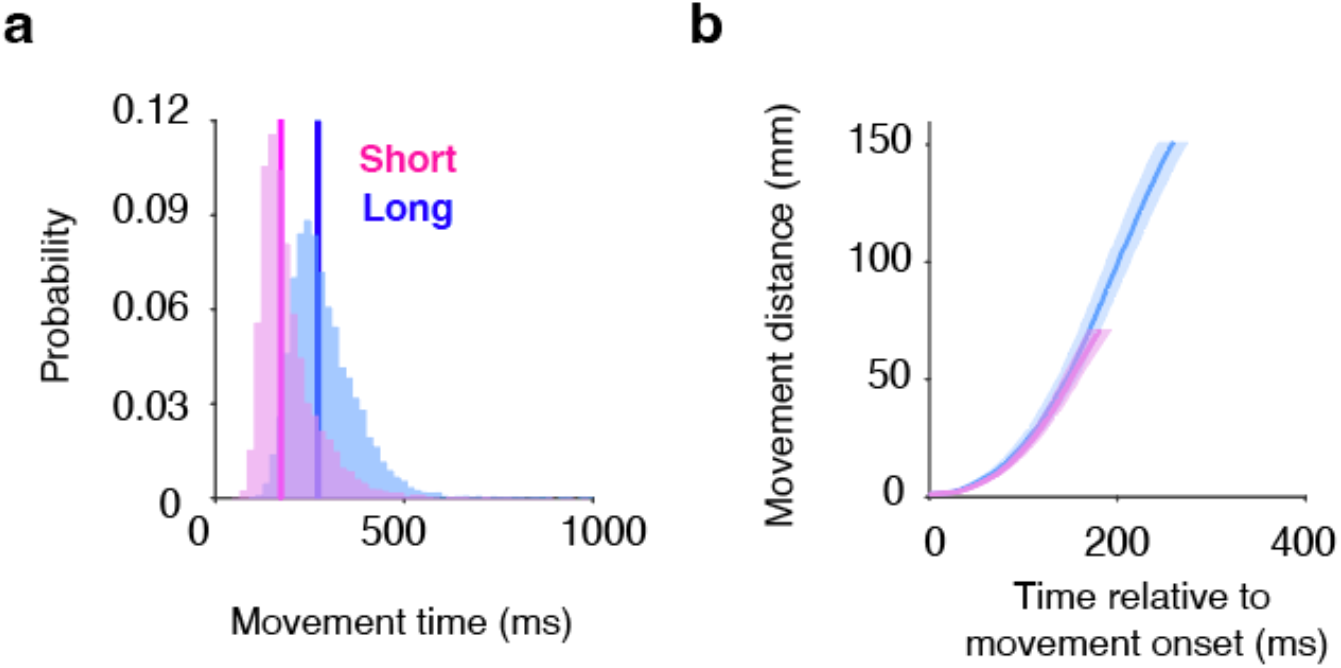
Increasing movement distance was effective manipulation for increasing movement duration in Experiment 3. **a**, Movement duration distributions for the short and long conditions. Thick vertical lines indicate group medians. **b**, Average hand distance as a function of time, relative to movement onset.

**Figure S7.**
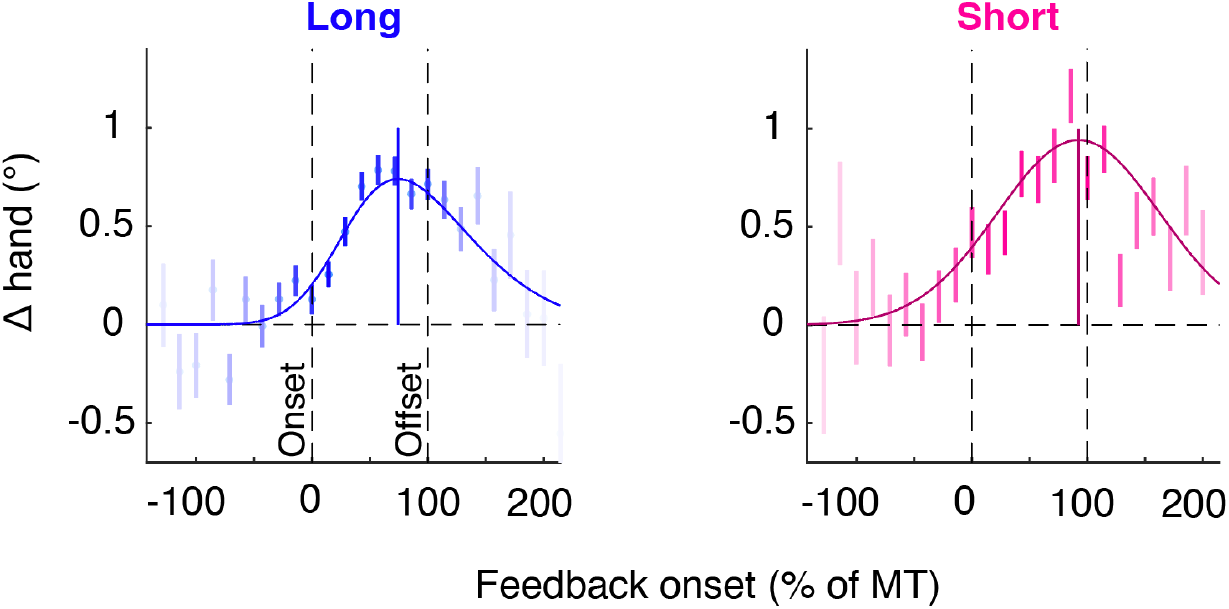
Trial-by-trial motor correction plotted as a function of feedback of normalized movement time (based on when hand reached target amplitude). The feedback onset time for each trial was aligned with movement onset and then divided by the movement duration of that trial. Bin size is 15% and the darkness of the dot indicates the relative number of samples in each bin. Colored curve indicates the best-fitted skewed Gaussian, with the colored vertical line marking the peak of the function.

**Table S2:**
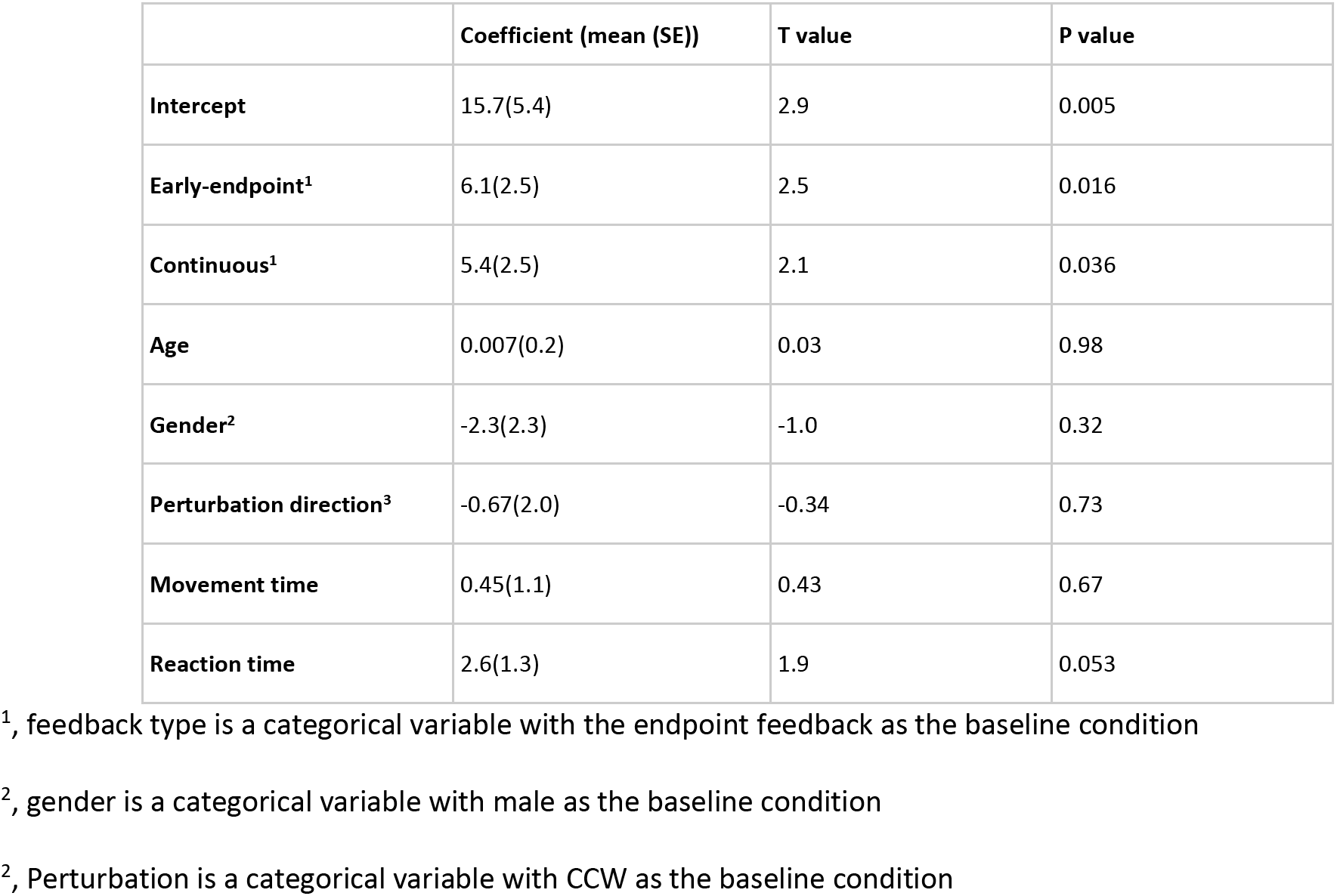
Linear regression results for Experiment 1

